# Walking with increased step length variability increases the metabolic cost of walking

**DOI:** 10.1101/2024.05.28.596299

**Authors:** Adam B. Grimmitt, Maeve E. Whelan, Douglas N. Martini, Wouter Hoogkamer

## Abstract

Older adults and neurological populations tend to walk with slower speeds, more gait variability, and a higher metabolic cost. This higher metabolic cost could be related to their increased gait variability, but this relationship is still unclear. The purpose of this study was to determine how increased step length variability affects the metabolic cost of waking. Eighteen healthy young adults completed a set of 5-minute trials of treadmill walking at 1.20 m/s while we manipulated their step length variability. Illuminated rectangles were projected onto the surface of a treadmill to cue step length variabilities of 0, 5 and 10% (coefficient of variation). Actual step lengths and their variability were tracked with reflective markers on the feet, while metabolic cost was measured using indirect calorimetry. Changes in metabolic cost across habitual walking (no projections) and the three variability conditions were analyzed using a linear mixed effects model. Metabolic power was largest in the 10% condition (4.30 ± 0.23 W/kg) compared to 0% (4.16 ± 0.18 W/kg) and habitual (3.98 ± 0.25 W/kg). The participant’s actual step length variability did not match projected conditions for 0% (3.10%) and 10% (7.03%). For every 1% increase in step length variability, there is an 0.7% increase in metabolic cost. Our results demonstrate an association between the metabolic cost of walking and gait step length variability. This suggests that increased gait variability contributes to a portion of the increased cost of walking seen in older adults and neurological populations.

**Summary Statement:** For every 1% increase in step length variability, there is an 0.7% increase in the metabolic cost of walking.

## INTRODUCTION

With age and increased neurological impairment, people walk slower (Bohannon, 1997; Steffen et al., 2002), with shorter steps (Osoba et al., 2019), greater gait variability (Owings & Grabiner, 2004) and increased metabolic energy demands (Christiansen et al., 2009; Martin et al., 1992; Zamparo et al., 1995). Increased metabolic cost of walking has been mentioned as a potential risk factor for reduced gait speed and mobility in older individuals (Schrack et al., 2012). Reduced gait speed is associated with mortality (Newman et al., 2006; Studenski et al., 2011), cardiovascular disease, and other adverse effects (Abellan van Kan et al., 2009; Szanton et al., 2021). As outlined by Boyer et al., (2023) “… the significant societal, economic, and personal burdens associated with mobility limitations highlight the importance of understanding the mechanisms for increased metabolic cost of walking…”. While the association between reduced gait speed and increased metabolic cost is well studied (but not yet fully understood; Boyer et al., 2023), less is known about the potential association between other gait changes, such as increased gait variability, and the metabolic cost of walking.

Increasing stride frequency from habitual (+15 strides / minute) increases metabolic cost by approximately 19% during walking (Holt et al., 1991). As step frequency increases, there is a greater cost of moving the legs (Doke et al., 2005). Decreasing stride frequency from habitual (– 15 strides / minute) increases metabolic cost by almost 30% (Holt et al., 1991). This is likely due to the increased metabolic cost of redirecting the center of mass between steps (Donelan et al., 2002). Indeed, people habitually self-select a step frequency which minimizes metabolic cost (Holt et al., 1995).

However, people do not walk with consistent step lengths in daily life because walking often occurs in short bouts (Seethapathi & Srinivasan, 2015), at variable speeds, and not on level ground (Kowalsky et al., 2021; Voloshina et al., 2013). Varying step length can also be helpful when encountering obstacles (Patla et al., 1991) or maintaining stability (Young & Dingwell, 2012). Even for a long walking bout, at a constant speed, on level ground, with one’s optimal average step length, there will be small variabilities in step length around this average - a mix of steps that are both shorter and longer than optimal (Owings & Grabiner, 2004), due to intrinsic sensory and neuromotor noise (Dean et al., 2007; Dingwell et al., 2017). Added variability can be expected to be more metabolically costly (O’Connor et al., 2012), as each step deviates from optimal step parameters (Holt et al., 1995).

Few studies have tried to quantify the impact of step length variability on metabolic cost (O’Connor et al., 2012; Rock et al., 2018). O’Connor et al. (2012) used virtual visual flow field perturbations to increase gait variability and evaluate metabolic cost. At a constant speed of 1.25 m/s, high frequency medio-lateral rotations of the virtual visual flow field induced the largest increase in metabolic cost (5.9%). Participants walked with increased step length variability in this condition, but the metabolic increase was most strongly coupled with increased step width variability. More recently, Rock et al. (2018) evaluated metabolic cost of transport and step length variability across a range of walking speeds. Although speed had the greatest impact on metabolic cost, at a constant speed of 1.25 m/s, each 1% increase in step length variability was also associated with a 5.9% increase in metabolic cost. In these studies, the observed increases in metabolic cost with increased step length variability could not be separated from changes in step width variability or walking speed, respectively, so the isolated effects of increased step length variability on the metabolic cost of walking are still unknown.

Therefore, we set out to quantify the isolated effects of increased step length variability on the metabolic cost of walking. We used visual cues to increase step length variability by projecting illuminated rectangles (stepping stones) on a treadmill progressing at the speed of the belt (Hollands et al., 1995; Hoogkamer et al., 2015; Roerdink et al., 2009; Van Ooijen et al., 2015). The stepping stones were spaced to the participant’s habitual step length, with increasing levels of step length variability per condition. By directly targeting step length variability, using projected stepping stones, we were able to evaluate the metabolic cost of walking with increased step length variability independent from other gait changes that may arise when using perturbations that indirectly affect step length variability. We hypothesized that increases in step length variability would increase the metabolic cost of walking.

## MATERIALS AND METHODS

### Participants

Eighteen healthy young adults (7F; 24.4 ± 3.7 years; 171.2 ± 17.2 cm; 70.5 ± 13.3 kg) completed this study. Eligible participants were between the ages of 18 and 45 years old, had not experienced lower extremity injuries or surgery within the past six months, and were free of any existing orthopedic, cardiovascular or neuromuscular conditions. Written informed consent was obtained from each participant prior to the study. All procedures were approved by the Institutional Review Board at the University of Massachusetts Amherst (#3002).

### Procedures

We provided each participant with a pair of standardized shoes in their size (Speed Sutamina, PUMA SE, Herzogenaurach, Germany). We placed retroreflective markers on each foot at the fifth metatarsal head and calcaneus, and a four-marker cluster on the sacrum.

Participants first walked at a speed of 1.20 m/s (Das Gupta et al., 2019) for five minutes to familiarize themselves with the dual-belt treadmill (Bertec, Columbus, OH, USA) and the indirect calorimetry mouthpiece. Next, they walked for three minutes at 1.20 m/s for which we evaluated habitual step length during the final 30 seconds. An eight-camera Miqus system (Qualisys, Gothenburg, Sweden) recorded kinematic data at 100 Hz. We used kinematic data from the left and right calcaneus to determine habitual step length using a custom Matlab script (The MathWorks, Natick, MA, USA). Step length for each leg was calculated as the distance between the anterior and posterior positions of the ipsilateral and contralateral calcaneus markers, respectively, during the maximum anterior position of the calcaneus marker for each step (Desailly et al., 2009). Mean step length and standard deviation were used to determine the coefficient of variation, i.e., the mean step length divided by the standard deviation for each foot, before averaging across feet.

Experimental conditions were: no projections (NP), 0%, 5% and 10% variability. Throughout we consider step length variability in terms of the coefficient of variation of the step length during a trial. At 0% variability, we projected stones with no variability in step lengths, whereas for the 10% condition we projected stones with 10% variability in step lengths across the entire trial. In a block randomized order, participants completed all four experimental conditions, before completing them again in reverse, for a total of eight trials (e.g., NP, 10%, 0%, 5%, 5%, 0%, 10%, NP).

The projected stepping stones were generated using a custom Matlab script. A vector of step lengths for the entire trial was created based on the participant’s habitual step length, treadmill speed (1.20 m/s), and trial duration (5 minutes). A vector of corresponding step length perturbations was created using the *randn* function in Matlab, which generates a list of normally distributed random numbers with a mean of 0 and variance of 1. This step length perturbation vector was scaled by the desired step length variability (0, 5 or 10% coefficient of variation) and the habitual step length. A larger coefficient of variation increases the frequency of perturbed steps relative to unperturbed, and of larger perturbations relative to smaller. To ensure that differences in step length were perceivable by the participants, we discretized the perturbations into bins that were 5% multiples of the participant’s habitual step length. We limited perturbations to a maximum of 15% in either direction, to prevent the possibility of unrealistic perturbations at the tails of the normal distribution. Perturbations that fell within half of the discretization interval above or below a perturbation target were reassigned to the target value and applied. Thus, participants encountered distances that matched the habitual step length most of the time, with less frequent longer and shorter perturbed steps (Fig. 1A).

**Figure 1.**
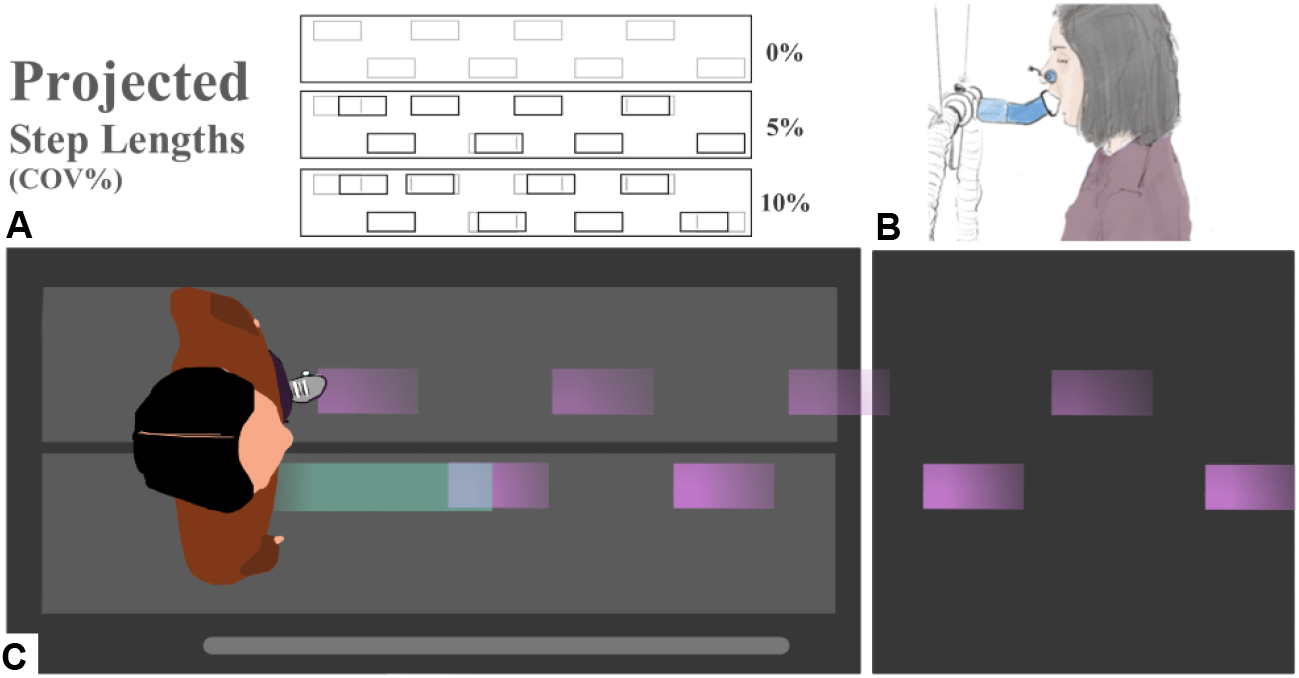
(A) Diagram of example step length conditions for 0, 5, and 10% variability. Grey rectangles are stepping stone targets projected at a participant’s specific habitual step length. Black rectangles are the lengthwise deviations of perturbed stones generated randomly from discretized bins. (B) Illustration of indirect calorimetry including pitched 45-degree extension used to open participant field-of-view. (C) Top-down visual of a participant targeting projected stepping stones (purple; moving from right to the left) for 0% variability.

We used an expired-gas analysis system (True One 2400, Parvo Medics, Salt Lake City, UT, USA) to measure metabolic cost across the four experimental walking conditions. We added a 3D printed extension to the three-way valve mouthpiece (Hans Rudolf Mouthpiece, Shawnee, KS, USA) that pitched forward (5 cm) and up (5 cm) at a 45-degree angle (Fig. 1B). The participants were then able to see the approaching stepping stones. Each trial was five minutes long and data across the last two minutes of each trial was used to evaluate metabolic power. We calculated metabolic power (W/kg) using oxygen uptake, carbon-dioxide production, and the Péronnet and Massicotte equation (Kipp et al., 2018; Péronnet & Massicotte, 1991). Metabolic power was averaged across the two trials for each condition (Barrons et al., 2024).

### Statistical Analysis

We used linear mixed effects models to evaluate changes in metabolic cost with increasing step length variability. We used the actual step length variability that participants walked with for each condition, rather than the projected step length variability for that condition. All statistical analysis was performed in R studio (4.2.2) with a linear mixed effects regression (Wilkinson et al., 2023) using lme4 (1.1 - 32) and sjstats (0.18.2) packages. A paired t-test was used to compare metabolic power and step length variabilities between NP and 0% conditions. A linear model was used to investigate the impact of step length variability on metabolic power during walking. Random intercepts were adjusted for each participant using equation 1 where COV is the coefficient of variation of the step lengths during a trial.

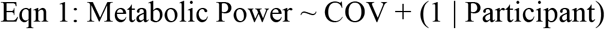

Metabolic power was then evaluated using the mixed effect model with a random intercept for each participant and an independent slope corresponding to the participant specific metabolic power response using equation 2.

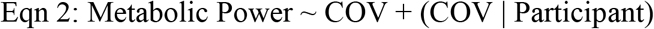

We used a likelihood ratio to test for significance between a model excluding step length variability and an alternative model that did not. Chi-squared (χ^2^) and significance values are reported. For all statistical tests, significance was set at an alpha level of 0.05.

## RESULTS

Metabolic power and step length variability were different between NP and 0% conditions (3.98 ± 0.25 vs. 4.16 ± 0.18 W/kg; p = 0.0002 and 2.42 ± 0.55% vs. 3.10 ± 0.58%; p = 0.001, respectively; Figure 2). Thus, when pooling the true step length variability across conditions, we excluded the NP condition (i.e., we included only data from the 0%, 5% and 10% step length variability trials in our model). Overall, metabolic power increased with increasing step length variability (χ^2^ = 9.41; p = 0.002; Fig. 3). For every 1% unit increase in step length variability, there was an increase in metabolic power of 0.03 ± 0.01 W/kg, or 0.7%.

**Figure 2.**
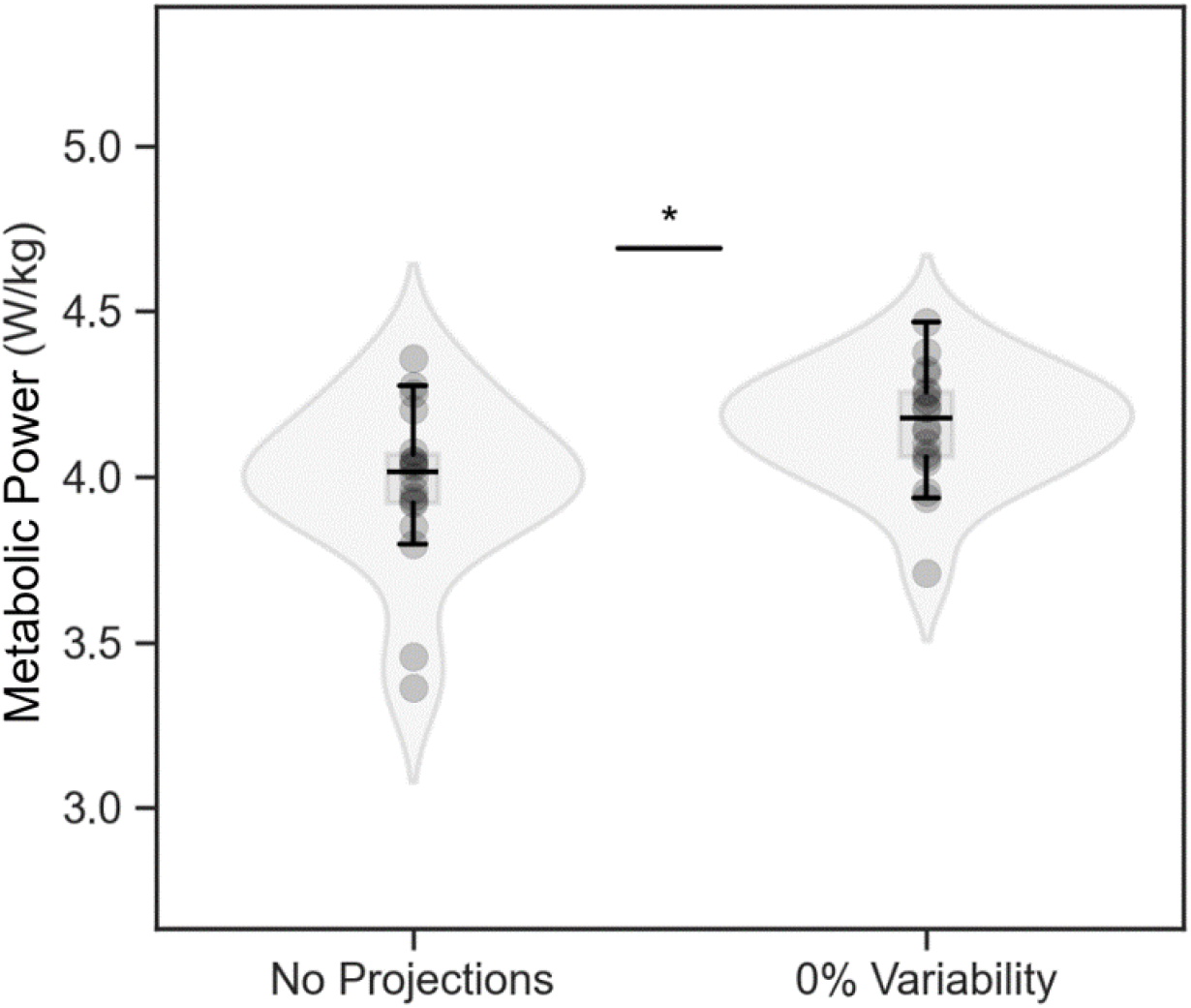
Metabolic power (n = 18) was lower for walking without any projected stepping stones than for walking over stepping stones projected without any step length variability. Values are mean ± s.d., * *p* < *0.05*

**Figure 3:**
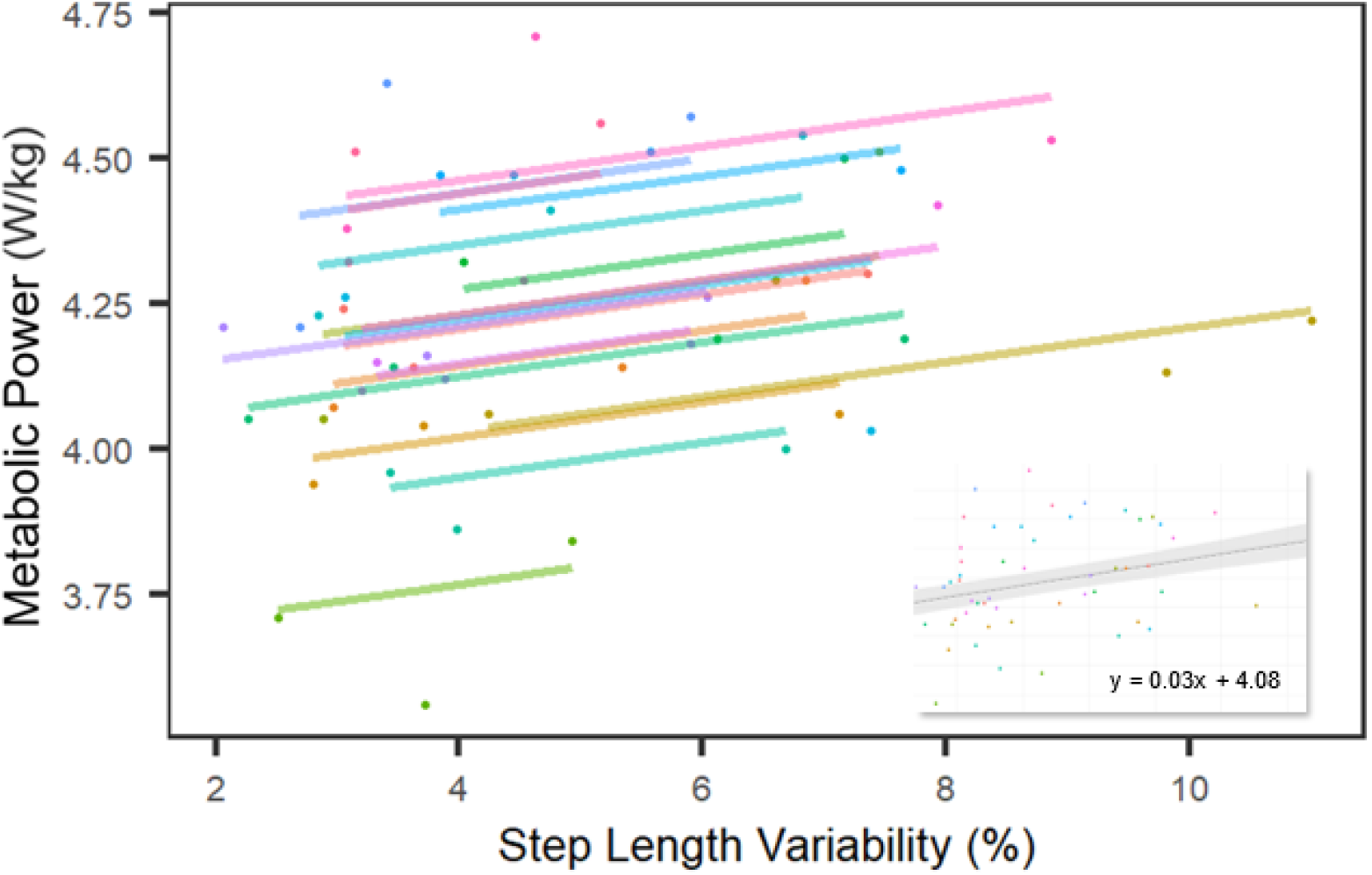
For every percentage increase in step length variability, there is a 0.03 W/kg (0.7%) increase in metabolic power, averaged between conditions and across participants (inset). Linear mixed effects models identified linear trends for each subject (n =18; colored lines) across conditions.

## DISCUSSION

In this study, we quantified the isolated effects of increased step length variability on the metabolic cost of walking. In line with our hypothesis, increases in step length variability resulted in increases in metabolic cost. Additionally, we observed a difference in metabolic cost between walking without projections and with projections with no variability. Our results are similar, but of smaller magnitude than observations from studies that indirectly increased step length variability (O’Connor et al., 2012; Rock et al., 2018).

Our findings suggest a modest contribution of step length variability to the metabolic cost of walking (0.7% increase in metabolic power per 1% increase in step length variability; Fig. 3) at a common walking speed of 1.20 m/s. At this speed, every percent change in step length variability increases metabolic cost by 0.03 W/kg. This increase is almost five times smaller than the 0.14 W/kg increase in metabolic cost, for every 1% increase in variability, modeled by Rock et al (2018). This difference is likely related to our direct manipulation of step length variability at a single, constant speed, eliminating the metabolic penalty of walking slower than preferred (Ralston, 1958). Additionally, the work of O’Connor et al. (2012), reported greater step width variability (+65%), mean step width (+19%) and an increased metabolic cost (5.9%) while walking with virtual visual flow perturbations (high frequency medio-lateral rotations). This suggests that step width variability has a larger effect on metabolic cost than step length variability. Indeed, walking with increased step width comes at a metabolic cost (Donelan et al., 2001).

Two mechanisms can be expected to play a role in elevating the metabolic cost of walking with increased step length variability. First, adjusting to a closer stepping stone (i.e., reducing step length), increases the metabolic cost of moving limbs at a higher rate (Doke et al., 2005). Second, an adjustment to a farther stepping stone (i.e., increasing step length) increases the cost of lifting the center of mass over the point of collision (Donelan et al., 2001). An additional possible mechanism, that relates to the need to step accurately on each stone, is an increase in muscle co-contraction, commonly observed with increased accuracy demands (Gribble et al., 2003). While some of the increased metabolic cost of walking could be attributed to increased accuracy demands, an additional portion could be related to the cost of consistently regulating steps across projected stones, degrading the contribution of passive dynamics to walking (Wezenberg et al., 2011). These additional mechanisms could help explain the differences in metabolic cost between walking with no projections (habitual) and with 0% step length variability projections that closely matched habitual gait characteristics (Fig. 2).

Beyond slower gait with shorter steps and longer double support phases, older adults have more gait variability and increased metabolic energy demands (Boyer et al., 2023; Schrack et al., 2012). The larger gait variability in older adults and in those with neurological impairments could contribute to their increased metabolic cost of walking as compared to young (e.g., +8%; Martin et al., 1992) or neurologically healthy adults (+17-170%; Compagnat et al., 2020; Jeng et al., 2020; Rooney et al., 2022). At matched speeds, older adults walk with higher step length variability than younger adults (Almarwani et al., 2016; Kang & Dingwell, 2008), more so if the older adult has mobility impairments, with a reported 2.7% increase in step length variability (James et al., 2020). Our data suggest that a 2.7% increase in step length variability would increase the metabolic cost of walking by 1.7%. At their preferred speeds, adults with neurological impairments have been reported to walk with step length variabilities that are approximately 2.5%, 3.0%, and 6.0% larger than neurologically healthy controls, for Parkinson’s disease, Multiple Sclerosis, and Cerebellar Ataxia, respectively (Buckley et al., 2018; Noh et al., 2020; Roemmich et al., 2012; Socie et al., 2013). Our data suggests that these increases in step length variability will increase the metabolic cost of walking by 1.8, 2.1 and 4.2%, respectively. The metabolic cost of walking with increased step length variability only contributes a small part to the total increased metabolic cost (17– 170%) of walking observed across these populations (Compagnat et al., 2020; Jeng et al., 2020; Martin et al., 1992; Rooney et al., 2022).

### Limitations and future directions

To ensure that the step length variations were perceivable to participants, we binned step lengths into 5% multiples, not exceeding 15% of their habitual step length. Normal walking does not contain such discretized step length variability. Although the perturbed conditions do not perfectly reflect real-world gait variability, they provide an upper limit for the increases in metabolic cost in young healthy individuals. Most participants were unable to exactly match the discretized step length variabilities that were projected; while the use of 5% bins is different than real-world gait variability, participants still had trouble maintaining accuracy across the stepping stones without additional feedback. We were limited in identifying how participants altered their step length. In future research it could be insightful to evaluate whether participants took larger or shorter steps when aiming for the stone targets, as well as the frequency and magnitude of their errors. This would not change the relationship between step length variability and metabolic cost, but it could help in relating task performance to step length variability within a condition. While few participants were able to walk with 5 and 10% step length variability, 0% step length variability was unachievable – the lowest value for a single participant was 2%. This is in line with observations that walking inherently contains some variability in step length (Collins & Kuo, 2013) which might also be distinct between participants.

Participants did not receive feedback on how well they were performing during a trial. Feedback could have improved task performance (Shull et al., 2014) to better align with our variability conditions. To investigate whether co-contraction, to maintain accuracy (Gribble et al., 2003), contributes to the increased metabolic cost of walking with variable step lengths, future studies should quantify foot placement accuracy and muscle activity across similar virtual projections. Finally, to clarify the relationship between variability and gait deficiencies for populations already experiencing increased step length variability, further investigations should be conducted with these populations specifically.

### Conclusion

Older adults and those with neurological conditions walk with greater step length variability and increased metabolic cost. In a population of healthy young adults, we found that metabolic cost increases by approximately 0.7% (0.03 W/kg) for every 1% increase in step length variability. Although most of our participants were unable to exactly match the projected conditions, our virtual visual perturbation successfully increased step length variability and metabolic cost across step variability conditions. The metabolic cost per unit of variability was smaller than reported in previous work, indicating that step length variability plays a modest, albeit significant role in the metabolic cost of walking.

## Acknowledgments

We acknowledge our volunteers from the University of Massachusetts Amherst who assisted in piloting and data collection, the Kinesiology staff at the University of Massachusetts Amherst, Dr. Mark Price for his assistance in projecting a pathway and Dr. Dan Feeney for helping us tactfully compare humans walking over it.

## Competing interests

The authors declare no competing or financial interests.

## Author contributions

Conceptualization: M.E.W., W.H.; Methodology: A.B.G., M.E.W., W.H. Formal analysis: A.B.G., M.E.W., W.H. Investigation: A.B.G., M.E.W.; Resources: A.B.G., W.H.; Data curation: A.B.G., M.E.W., W.H; Writing - original draft: A.B.G., M.E.W., WH; Writing - review & editing: A.B.G., D.N.M., W.H; Visualization: A.B.G; Supervision: D.N.M., W.H.; Project administration: W.H.; Funding acquisition: D.N.M., W.H

## Funding

This work was supported by the National Institute of Health (R21AG075489)

## Notes

### Competing Interest Statement

The authors have declared no competing interest.

